# Large vision model framework for automated *C. elegans* analysis: From static morphometry to dynamic neural activity

**DOI:** 10.1101/2025.08.18.670800

**Authors:** Aurélie Guisnet, Michael Hendricks

## Abstract

Quantitative phenotyping of *Caenorhabditis elegans* is essential across numerous fields, yet data extraction remains a significant analytical bottleneck. Traditional segmentation methods, typically reliant on pixel-intensity thresholding, are highly sensitive to variations in imaging conditions and often fail in the presence of noise, overlaps, or uneven illumination. These failures necessitate meticulous experimental setups, expensive hardware, or extensive manual curation, which reduces throughput and introduces bias. Here, we introduce TWARDIS (Tools for Worm Automated Recognition & Dynamic Imaging System), a modular, Python-based analysis suite that leverages large foundation vision models, specifically the Segment Anything Models (SAM and SAM2) and a fine-tuned vision transformer classifier, to overcome these limitations. We demonstrate the versatility and robustness of our AI compound system across diverse modalities. For static morphological analysis, TWARDIS successfully resolved overlapping worms in noisy images without human intervention, showing a 0.999 correlation with manual segmentation. In behavioral assays (swimming and crawling), the pipeline enabled high-definition postural analysis even in low-resolution, wide-field recordings where the worm occupied only ∼0.25% of the field of view, accurately resolving complex postures without frame rejection. Finally, when applied to calcium imaging of semi-restricted animals, TWARDIS provided precise, frame-by-frame segmentation of neural compartments, reducing the artificial signal flattening common in traditional region-of-interest-based approaches and enabling the extraction of biologically accurate, absolute head positions. The system’s hardware-scalable architecture and modular design ensure both current accessibility and future improvements without restructuring. By automating the most time-consuming aspects of image analysis, TWARDIS removes critical bottlenecks and tradeoffs in *C. elegans* research, enabling researchers to focus on biological questions rather than technical image processing challenges.

**Author Summary:** The small roundworm *Caenorhabditis elegans* is widely used by scientists to study fundamental biological questions, such as aging and how the brain works. A crucial part of this research involves analyzing images and videos to measure the worm’s shape, movement, and neural activity. However, extracting accurate data is a major challenge. Traditional software tools often fail if the lighting is uneven, the image is noisy, or worms overlap. This forces researchers to spend countless hours manually correcting errors or investing in expensive, specialized equipment. To overcome this bottleneck, we developed TWARDIS (Tools for Worm Automated Recognition & Dynamic Imaging System). Our system utilizes recent advances in Artificial Intelligence, harnessing powerful, generalized AI models to automatically and accurately identify the worms, even in challenging images. We demonstrated that TWARDIS reliably analyzes complex behaviors and neural activity without human intervention. Our approach removes the traditional trade-off between data quality and equipment cost, enabling high-precision analysis using simple setups. By automating the most tedious parts of image analysis, TWARDIS allows scientists to focus on biological discovery rather than technical hurdles.

## Introduction

The nematode *Caenorhabditis elegans* has long been established as a leading model organism in biological research, owing to its genetic tractability, conserved biological pathways, short life cycle, and optical transparency. These attributes make it invaluable for studies spanning aging, development, neurobiology, and drug discovery. A cornerstone of *C. elegans* studies is the ability to quantitatively phenotype the organism, which involves, for example, accurately measuring morphology, characterizing complex behaviors, or tracking neural activity. While imaging technology has become increasingly accessible, the extraction of meaningful, high quality data from these images remains a significant time consuming analytical bottleneck.

The first and most critical step in virtually all image-based phenotyping pipelines is segmentation: the process of accurately delineating the organism or structures of interest from the background. Traditionally, this has relied heavily on pixel intensity thresholding and basic feature detection algorithms [1,2]. However, these methods are notoriously fragile and highly sensitive to experimental variability. Imperfections common in routine data acquisition, such as uneven illumination, variations in focus, background debris, or overlapping individuals, frequently cause threshold-based methods to fail, yielding fragmented or erroneous segmentations [3,4].

To compensate for these failures, researchers often invest significant effort in optimizing imaging conditions, utilize expensive high-resolution hardware, compromise on limited feature extraction or resort to extensive manual curation [5]. In dynamic analyses, such as behavioral tracking or calcium imaging, poor segmentation frequently leads to the rejection of frames or the artificial flattening of signals, resulting in the loss of critical temporal data and potentially masking biological phenomena. Semi-automated tools often require user input to define regions of interest, adjust parameters or manually curate results, which increases processing time and introduces inter-observer variability and bias [2,6,7]. Furthermore, many existing automated tools require specialized computational environments or proprietary licenses (e.g., MATLAB, C++, or complex Docker configurations), limiting their accessibility.

Recent advancements in deep learning have shown promise in automating *C. elegans* analysis [3,8,9]. While powerful, many of these approaches required extensive, expertly annotated datasets tailored to specific imaging conditions and tasks, limiting their generalization across different experimental setups. A transformative development in computer vision has been the emergence of vision transformer-based foundation models, some of which are trained on extremely diverse, high quality datasets and can perform new tasks with minimal or no fine-tuning. Notably, the Segment Anything Model (SAM) [10] and its successor, SAM2 [11], offer unprecedented generalization capabilities for object segmentation, demonstrating remarkable robustness to image variability without customization. The success of foundation models in medical imaging, microscopy, and other biological applications suggests their potential to transform *C. elegans* research as well [12–14].

Here, we introduce TWARDIS (Tools for Worm Automated Recognition & Dynamic Imaging System), a modular, Python-based analysis suite that harnesses the power of these foundation vision models to overcome long-standing compromises in *C. elegans* phenotyping. TWARDIS provides automated, high-precision pipelines that require minimal user input and is resilient to the imperfect conditions typical of routine laboratory imaging. We demonstrate the versatility of TWARDIS across four modalities of increasing complexity: multi-worm morphological feature extraction from static images; high-definition analysis of swimming behavior in liquid; tracking and postural analysis of crawling worms in low-resolution, wide-field environments; and precise segmentation of neural compartments and head kinematics during calcium imaging. Across all applications, TWARDIS reduces the need for manual intervention, addressing the fundamental tension between the high-throughput nature of *C. elegans* experiments and the labor-intensive nature of image analysis. The system’s pure Python implementation and hardware scalability ensures accessibility across users. As foundation models continue to improve, TWARDIS pipelines will inherit these advances without requiring fundamental restructuring, providing a sustainable solution for evolving needs. This work demonstrates how leveraging generalized AI capabilities through a compound system, rather than developing increasingly specialized solutions, can eliminate longstanding technical bottlenecks, improve community adoption and accelerate biological discovery.

## Results

### Static multi-worm morphological feature extraction

We first integrated the SAM vision model with our fine-tuned worm classifier to automatically extract morphological features from static images of worms. After initial image segmentation with the SAM model, we fine-tuned a vision transformer classifier to automatically select worm segments (Fig 1A). We tested this TWARDIS pipeline across diverse worm life stages (L1 larvae to day 6 adults). We successfully segmented images and used our fine-tuned classifier to select worm segments. We obtained perfect classification of segments at all life stages in a single forward pass with no human intervention, demonstrating remarkable resilience to common imaging challenges such as variability in lighting, focus, noise artifacts and sample preparation. Traditional binary thresholding methods, and even more modern approaches, are sensitive to image imperfections: for *C. elegans* images, they often fail on out of focus elements, worm overlaps, sharp pharynx contrast, or noisy images, requiring either meticulous experimental setups for near perfect data acquisition conditions with manual or automated curation to discard invalid samples [2–4]. Visual comparisons in Figs 1B–F exemplify these advantages: raw images with overlapping worms and noise yield fragmented or erroneous outputs via thresholding, while our implementation produces clean, accurate masks without requiring any manual input, such as clicking on objects or defining regions of interest [6,15]. We tested our worm classifier on a second set of images collected on another recording set up and obtained zero false positives and less than 5% of false negatives showing that over-fitting on the optical profile of our training set was minimal (Figs 1D–F).

**Fig 1.**
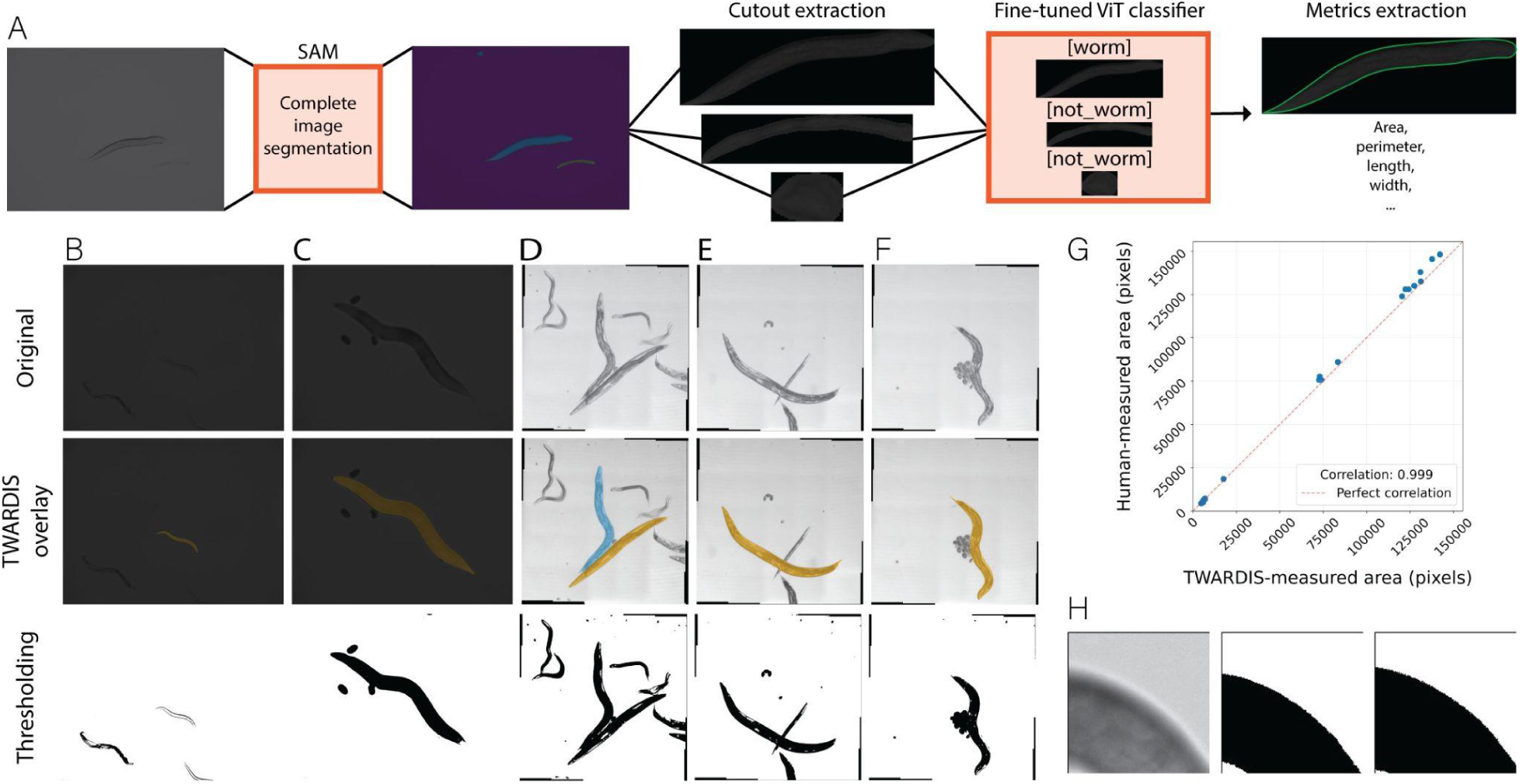
Multi-worm morphological feature extraction in static images. (A) Multi-worm static image feature extraction workflow. Each image is first passed through SAM for a complete segmentation. Cutouts are extracted for each identified mask that is not touching the sides of the image. The cutouts are classified as “worm” or “not worm” by our fine-tuned ViT classifier. Metrics are extracted for each identified worm mask. (B–F) Example comparisons of worm segmentation by thresholding versus TWARDIS age-based extraction. A–B samples from the first dataset. C–E samples from the second dataset. (G) Area in pixels of randomly selected worms measured by human manual segmentation versus TWARDIS-based segmentation. n = 21. (H) Example discrepancy in edge placement when thresholding on a blurry image.

From these segmented masks, we extracted key morphological features, including area, perimeter, length, and width along the entire worm length with minimal additional computational effort. When compared to a manual segmentation, the correlation with our TWARDIS implementation was 0.999 (Fig 1G). No images from our dataset could be correctly segmented by using only traditional thresholding, showing that our method dramatically reduces the need for pre-acquisition care or post-analysis handling time (Figs 1B–F).

This fully automated approach not only eliminates user intervention but also mitigates inter-observer variability and bias inherent in manual annotation [16,17]. For instance, multiple researchers outlining the same worm contours, either manually or by thresholding, will inevitably have discrepancies in edge placement which in turn can lead to variability in measured features (Fig 1H). In contrast, our method ensures a consistent approach across users and datasets. Most notably, our automation makes processing time irrelevant. By being fully automated, parallelizable and both CPU and GPU compatible, this TWARDIS workflow can be used to process any amount of data at a speed that scales with available hardware.

### Quantitative analysis of swimming behavior

We applied our approach to analyze video recordings of *C. elegans* swimming in liquid droplets (Fig 2A). This modality presents new challenges, including self-touching postures (extreme O- or 6-shapes), faint outlines, dynamic noise artifacts from light refraction, background fluctuations and moving extraneous particles. Current swimming analyses rely on thresholding and pixel differences for tracking which is less robust to imperfect conditions resulting in frame rejection or manual intervention to maintain accuracy [7,18]. In contrast, our TWARDIS implementation resolved these issues without any frame exclusions or manual tuning, producing high-definition masks that accurately captured worm shape even in a low-resolution space where the worm occupied only a very small portion of the total field of view (∼0.25%) (Fig 2F and S1 Movie). Our approach resulted in more complete body segmentations (Fig 2G–K).

**Fig 2.**
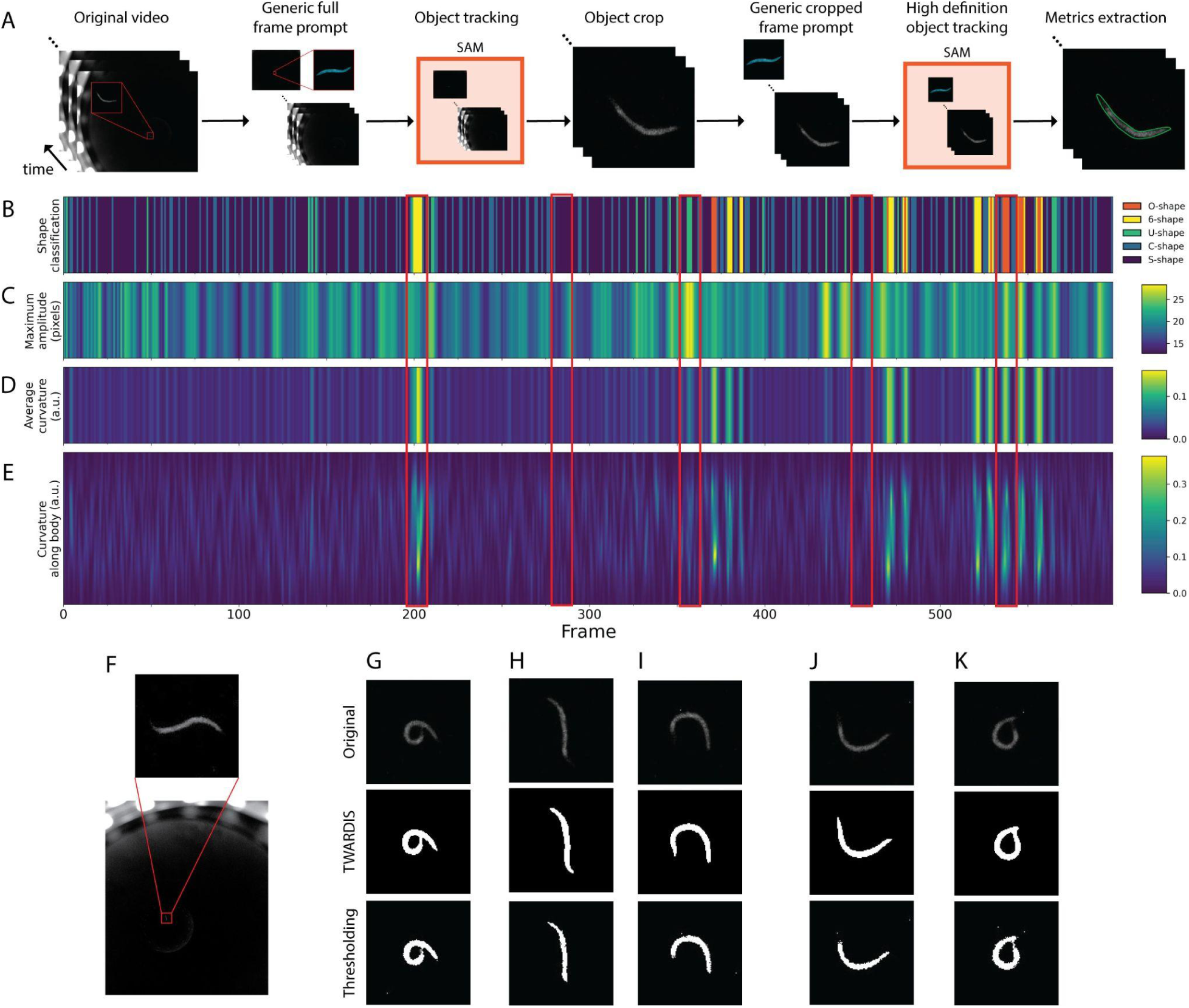
Quantitative analysis of swimming behavior. (A) Swimming behavior analysis workflow. First, a generic prompt frame is added to the video recording. Then, SAM is used to track the worm object through the recording. A cropped version of the video is created using the tracked worm. Another size-matched generic prompt frame is added to the cropped video and passed through SAM for a high-definition segmentation of the worm. Finally, metrics are extracted for each frame. Red picture-in-picture of worm shown for reference. (B–E) Selected time-series metrics of a 10 frames-per-second one minute recording of a wild-type worm swimming in a liquid droplet. (B) Per-frame shape classification based on amplitude and curvature. (C) Maximum bending amplitude per frame measured as the maximum perpendicular distance between the points along the worm’s body and a straight line connecting the head and tail. (D) Average body curvature per frame from E. (E) Gaussian-weighted curvature intensity of 100 interpolated points along the worm’s body per frame. (F) Example frame from recording representing the low space occupied by the worm within the total image field-of-view. (G–K) Original worm cutout with TWARDIS- and threshold-based segmentations for different frames along the recording. (G) Worms in a 6-shape are characterized by a small amplitude and very high curvature. (H) Worms in a mild S- or near straight shape are characterized by a very small amplitude and very small curvature. (I) Worms in a U-shape are characterized by a high amplitude and moderate curvature. (J) Worms in a wide C-shape are characterized by moderate amplitude and moderate curvature. (K) Worms in an O-shape are characterized by a high amplitude and high curvature.

From these refined masks, we extracted worm skeletons as described above, enabling the computation of a broad suite of additional behavioral features of interest with high fidelity such as curvature along the body, amplitude (perpendicular distance to the centerline) and wavelength. The superior foundation segmentation quality directly translated to more accurate skeleton extractions, eliminating the need to discard frames due to poor shape representation and allowing for continuous time-lapse metrics and precise timings of behaviors (Figs 2B–E). For example, deep 6-shaped bends can be distinguished as periods of very high curvature and moderate amplitude, whereas O-shaped bends have both high curvature and high amplitude (Figs 2G, 2K and S1 Movie). Using a single inexpensive camera, standard LED lighting and a simple set up, we achieved detailed feature extraction without the need for costly or careful preparations and removed the need for manual input.

### Single worm high-definition crawling tracking

We next extended our process to crawling videos of *C. elegans* on agar plates in unrestricted environments, where, like in our swimming recordings, worms occupied a small portion of the total field of view (Figs 3A and 3B). These simple recording conditions were also low in hardware cost and preparation time. Again, conventional thresholding methods typically struggle here, producing noisy or incomplete masks in the presence of self-touching postures or background artifacts, often leading to frame rejection, reliance on manual corrections or use of expensive recording equipment [19–21]. Our automated approach, however, resolved these challenges without any manual input or frame exclusions, achieving clean masks even in imperfect conditions (Fig 3B).

**Fig 3.**
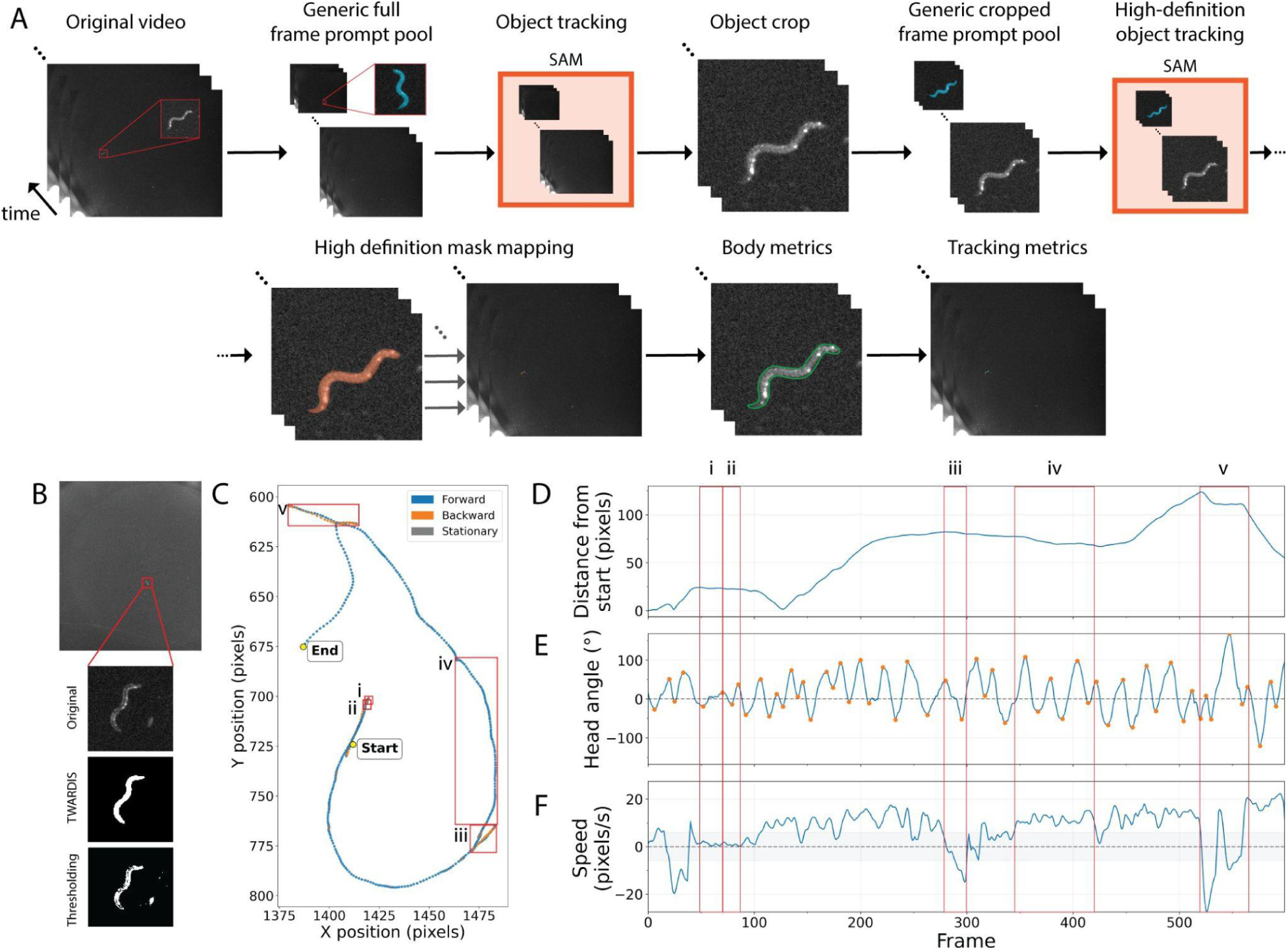
Single-worm tracking. (A) Crawling behavior analysis workflow. First, a generic prompt frame pool is added to the video recording. Then, SAM is used to track the worm object through the recording. A cropped version of the video is created using the tracked worm. Another size-matched generic prompt frame pool is added to the cropped video and passed through SAM for a high-definition segmentation of the worm. The high-definition segmented worm is mapped back to the original video. Finally, body metrics and path tracking metrics are extracted for each frame. Red picture-in-picture of worm shown for reference. (B) Example frame from recording representing the low space occupied by the worm within the total image field-of-view with an original worm cutout with TWARDIS- and threshold-based segmentations. (C–F) Selected time-series metrics of a 10 frames-per-second one minute recording of a wild-type worm crawling on an agar plate with food. (C) Worm path analysis color-coded by movement classification from (D) and where each point is a frame. Start and end points are labelled. X and Y-axes are centered (0,0) are the start point and units are pixels. (D) Euclidean distance from start point per frame in pixels. (E) Head bending angle relative to body in degrees per frame. 0 indicates a straight head relative to the body. Positive and negative values represent bends on either side, but ventral and dorsal sides are not distinguished. Points along the trace represent the detected peaks and troughs used for head bend metrics. (F) Worm speed by frame. Positive values represent forward motion, negative values represent backward motion, stationary motion is determined by absolute speed less than 0.6 pixel/second (shaded area). (C,Fi) Immobile stationary period. (C,Fii) Stationary period with head movement (C,Fiii) Reversal. (C,Fiv) Continuous forward crawl with a slightly curved path, distinguishable by head bends being deeper on one side. (C,Fv) Fast reversal followed by omega-turn (very deep head bend with sharp path direction change).

From these high-quality masks, we could extract the same body features as under swimming conditions in addition to a set of path features across frames, including, for instance, head bending angles, speed, and complete path tracking (Figs 3C–F). This enabled reliable body shape analysis alongside path tracking with complete frame inclusion, eliminating the common trade-off between centroid-only tracking more common with low-cost, basic systems versus full posture details which typically require more costly hardware [22]. By delivering high-resolution segmentation and a broad feature set from minimal hardware with no manual input, our TWARDIS approach bridges a gap between accessibility, precision and manual analysis time (S2 Movie).

### Segmentation of calcium imaging recordings of semi-restricted worms

Lastly, we applied our strategy to extract fluorescence information and head movements from calcium imaging recordings of semi-restricted *C. elegans* in microfluidic devices. Specifically, we tracked the fluorescence of the three axonal compartments of RIA (nrV, nrD and loop) in a microfluidic device where alternating streams of odors were presented to the animal while the head was free to move (Figs 4A and 4B) [23]. Our automated approach segmented each axonal compartment frame-by-frame, yielding precise masks regardless of structural change in the neuron. Using an expertly-annotated prompt pool of approximately 20 images to guide the model’s understanding of compartment boundaries, we obtained detailed and precise brightness measurements for calcium dynamics (Fig 4C and S3 Movie). Traditional segmentation tools include ImageJ, Matlab, or OpenCV software, where plugins or functions use image registration algorithms to align neuronal compartments across frames to extract fluorescence intensity by region-of-interest (ROI) selection [24]. However, these algorithms typically struggle with aligning structures that have inconsistent geometries across frames such as the three axonal compartments of the RIA interneuron. While this can be manually corrected, it significantly increases handling time. On the other hand, if these issues go undetected, errors are introduced in the data: in most cases, a sudden artificial drop in the fluorescence intensity caused by the axonal compartment going momentarily outside of the ROI results in the entire intensity trace to be flattened (Fig 4D). The loop compartment, being particularly dim at baseline, is particularly vulnerable from having an imperfect ROI selection. TWARDIS-based analysis resulted in loop fluorescence intensity traces having similar or greater mean absolute difference than Fiji-analyzed traces demonstrating less artificial flattening with our approach (Fig 4E).

**Fig 4.**
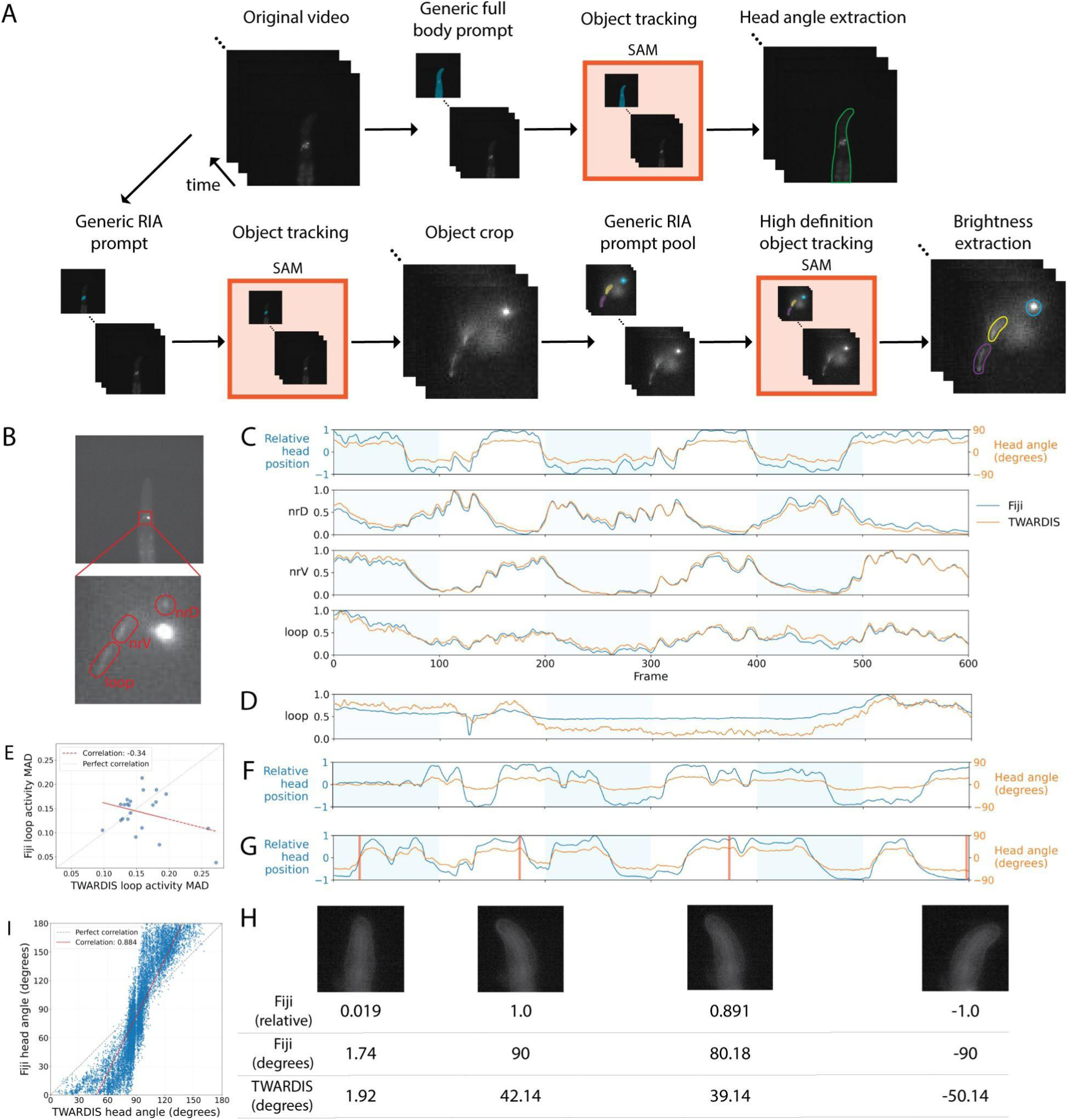
Segmentation of calcium imaging recordings of semi-restricted worms. (A) RIA calcium imaging analysis workflow. First, a generic prompt frame of the worm body is added to the video recording. SAM is used to track the worm object through the recording. The resulting worm masks are used to extract the head angles. Independently, a generic prompt frame of the RIA region is added to the video recording. SAM is used to track the RIA region through the recording. A cropped version of the video is created using the tracked RIA region. Another size-matched generic prompt frame pool of the three axonal compartments of RIA is added to the cropped video and passed through SAM for a high-definition segmentation of the compartments. The high-definition masks of the axonal compartments are used to extract brightness activity from the original recording. (B) Representative frame from calcium imaging recording with brightness enhanced for visualization. Blowup shows the three segmented compartments of the RIA interneuron whose fluorescence intensity through time are presented in (C). (C) Head bending and fluorescence time-series values of a 10 frames-per-second one minute recording of a wild-type worm in a microfluidic device expressing GCaMP3.3 in RIA. Blue traces represent Fiji-extracted data and orange TWARDIS-extracted data. Shaded areas represent periods where the positive odor is present. Top panel, Fiji-extracted relative head position (−1, 1) and TWARDIS-extracted head position (degrees relative to body). Second, third and fourth panel, nrD, nrV and loop normalized fluorescence intensity, respectively. (D) Illustrative normalized fluorescence intensity of the loop compartment where the Fiji-extracted data is artificially flattened. (E) Average absolute difference from mean for Fiji- versus TWARDIS-extracted data of the loop compartment. n = 22 recordings. (F-G) Illustrative traces where Fiji- and TWARDIS-extracted head positions are notably different. (H) Representative head positions from (G) with Fiji- and TWARDIS-extracted head position values. (I) Scatterplot of Fiji- versus TWARDIS-estimated head angles. Fiji estimations (−1, 1) are converted to the (−90, 90) range for comparison. 0 represents a perfectly straight head. +/− 90 degrees represent maximal bending. r = 0.884. n = 13,486 frames from 22 recordings.

Whole-body segmentation facilitated accurate head motion tracking via skeletonization and angle computation, providing more biologically relevant metrics. The classical approach is to estimate head position by using the maximum ellipse line of an ellipse fitted to a binarily thresholded version of the worm’s head. These measures are then normalized to the minimum and maximum for each recording to estimate head position on the 180 degrees of possible range. On the other hand, our approach yields the absolute head angle position relative to the midline of the body which provides much more accurate and relevant information for hypothesis testing (Figs 4C, 4F and 4G). In general, the ImageJ approach estimates deeper head bends than our TWARDIS approach with an overall correlation of 0.884 between the two approaches and of only 0.746 when direction of the head bend is ignored (Fig 4I and S1 Fig). These differences are particularly apparent in recordings where the worm’s head bends are minimal or restricted on one side where head angle inflation becomes most striking (Figs 4F–H and S3 Movie).

## Discussion

The quantitative analysis of *C. elegans* morphology, behavior, and neural activity is fundamental to its utility as a model organism. However, analytical pipelines in the field have often been constrained by a trade-off between precision, throughput, and accessibility. More precisely, traditional methods frequently require meticulous experimental conditions, expensive hardware, or significant manual intervention [2,5,21]. Furthermore, existing tools can be difficult to deploy, maintain or modify due to dependencies on proprietary software or complex programming environments (e.g., MATLAB, Java, C++, or Docker) [18,19,25–28]. This study introduces the first version of TWARDIS (Tools for Worm Automated Recognition & Dynamic Imaging System), a modular analysis suite implemented in Python that leverages the capabilities of foundation large vision models to deliver automated, high-precision phenotyping across diverse imaging modalities addressing key bottlenecks in researcher time spent on repetitive manual image analysis tasks and equipment requirements. We demonstrate robust and accurate segmentation across four distinct imaging modalities: static morphometry, swimming behavior, crawling locomotion, and calcium imaging, all while requiring minimal to no human intervention.

The core innovation of TWARDIS lies in addressing the fundamental limitation of existing analysis tools: the segmentation step. Virtually all current segmentation tools rely fundamentally on pixel-intensity thresholding or basic feature detection algorithms, even in more modern tools implementing neural network-based worm tracking [3,19,29–31]. These approaches are inherently fragile and highly sensitive to variations in lighting, focus, debris, and background noise. When these methods fail, they necessitate manual curation, frame rejection, or complex parameter tuning, creating analytical bottlenecks and introducing potential bias.

By utilizing the foundational Segment Anything Models (SAM) [10,11], TWARDIS moves beyond current limitations, demonstrating remarkable resilience to common imaging challenges. In our analysis of static morphological images, TWARDIS resolved overlapping worms and noisy backgrounds, across all life stages, where thresholding failed (Figs 1B–F). By integrating SAM with a fine-tuned Vision Transformer classifier, the pipeline generalized effectively across different recording setups. This automation not only accelerates analysis but fundamentally enhances reproducibility by eliminating the inter-observer variability inherent in manual annotation or threshold adjustment (Fig 1H) [16,17].

A significant advantage of the TWARDIS compound system is its ability to extract high-definition behavioral data from simple, low-cost recording setups. Detailed postural analysis has often been restricted to high-magnification expensive systems, while wide-field, low-resolution cheap recordings typically allow only centroid tracking [5]. TWARDIS bridges this gap, successfully analyzing both swimming and crawling behaviors even when the worm occupied only ∼0.25% of the total field of view (Figs 2F and 3B). The superior segmentation provided by the SAM model enabled the resolution of complex postures (Figs 2G–K). This precision allows for nuanced behavioral classification, distinguishing, for example, a deep 6-shaped bend (high curvature, moderate amplitude) from an O-shape (high curvature, high amplitude) (Figs 2G and 2K). Crucially, TWARDIS extracted similar metrics as other tools, but without discarding any frames or requiring manual inputs, preserving the temporal continuity essential for accurate analysis of behavioral sequences and dynamics (Figs 2B–E and 3C–F).

The application of TWARDIS to in vivo calcium imaging further highlights its precision in challenging dynamic contexts. Analyzing neural activity in semi-restricted animals, such as the three axonal compartments of the RIA interneuron, is difficult because the structures deform and shift during recording. Traditional ROI-based analysis, such as that implemented in Fiji, often fails to track these changes, leading to misalignment and artificial attenuation of the fluorescence signal when the structure momentarily exits the ROI or requiring extensive handling time for corrections (Fig 4D). TWARDIS, guided by prompt engineering, provides adaptive, frame-by-frame segmentation that captures the morphology of the neuronal compartments accurately. This resulted in more faithful representations of calcium dynamics, evidenced by reduced signal flattening frequent in the RIA loop compartment (Fig 4E).

Furthermore, TWARDIS significantly refines the measurement of behaviors during calcium imaging. We demonstrated that the standard method of fitting an ellipse to a thresholded head often inflates the estimation of head bends (Figs 4F–I). In contrast, TWARDIS leverages accurate whole-body segmentation to calculate the absolute head angle relative to the body midline. This provides a more interpretable and biologically relevant metric (Fig 4H), essential for precise correlation of neural activity and behavioral output.

The success of TWARDIS underscores the advantage of employing an AI compound system built upon a powerful, generalized foundation model. Because the Segment Anything Model is pre-trained on a vast dataset of remarkable quality, it provides exceptional, generalized segmentation capabilities without the need for extensive, modality-specific training required by many other deep learning approaches [3,6,8,29,32,33]. We harness this generalized power and use targeted strategies (a simple classifier and prompt engineering) with deterministic workflows to achieve high performance. This architecture makes the system easily transferable across different imaging settings and also modular, where both the pre-trained models and the deterministic workflows can be changed or improved. As newer versions of SAM and similar models achieve better performance with smaller computational footprints, TWARDIS will inherit these improvements without requiring fundamental restructuring. Requiring only a minimum of 4GB of RAM for the worm classifier, the TWARDIS workflow is compatible with both CPU and GPU architectures and parallelizable, thus scaling with available hardware. That said, this removal of the human-in-the-loop makes processing time largely irrelevant, as analyses can be run independently of user input (e.g., overnight).

While TWARDIS offers significant advancements, we recognize areas for future development. As of today, the early v0.1 TWARDIS scripts are available in a public repository (https://github.com/lillyguisnet/TWARDISv0.1), but we plan to release TWARDIS as a user-friendly, open-source Python package to facilitate widespread adoption and enhance accessibility in the future. The current worm classifier showed minor overfitting to our training dataset’s optical profile (less than 5% false negatives on a second setup; Figs 1B–F), which can be readily addressed by incorporating worm images available in public datasets to increase diversity. Currently, the pipeline is optimized for static multi-worm detection and recordings of single-worm swimming, single-worm crawling and RIA fluorescence imaging. Future iterations could aim to incorporate multi-worm tracking, multi-well swimming, diverse neuronal recording conditions or even freely moving calcium activity as we presented in [34]. Future versions should also leverage other recent computer vision approaches for skeleton extraction, multi-worm tracking and multi-worm overlap resolution [35–38].

In conclusion, TWARDIS provides a robust, precise, and automated framework for *C. elegans* phenotyping. By capitalizing on the strengths of foundation large vision models, our compound system overcomes the inherent limitations of traditional segmentation, enabling high-definition analysis even under imperfect imaging conditions and with low-cost recording equipment. This framework significantly reduces manual labor and improves data quality, paving the way for large-scale reproducible investigations that focus on experimental design and biological interpretation rather than manual image processing.

## Materials and Methods

### Static images acquisition and analysis

#### Developmental growth, first set up

Wild-type N2 *C. elegans* were grown at 20 C° and were fed OP50 *E. coli* bacteria as described in Guisnet et al. [39].Worms were timed by placing egg-laying adults on plates for 2 hours. These worms were moved to fresh plates through the experimental days as needed to keep them fed. Every 24 hours after that, worms were placed on 2% agarose pads in a 5 µL drop of 1M sodium azide to immobilize them. After 1 minute, the sodium azide was carefully absorbed with a Kimwipe. No coverslip was added to cover the worms as this caused them to be squished and deformed (particularly in width). Starting at day 3, focus was optimized for the head and tail as worms were too thick to be entirely in focus. Worms were imaged within 10 minutes of being immobilized. Grayscale images were taken with an Olympus XM10 CCD camera mounted on an Olympus MVX10 microscope using an MV PLAPO 2XC objective. TIF images were acquired with a 1376 x 1038 resolution and 10 ms exposure with the cellSens software (Evident).

#### Developmental growth, second set up

*C. elegans* (N2 wild-type strain) were grown at 20°C on nematode growth medium (NGM) agar plates seeded with *Escherichia coli* OP50. Synchronized worms were obtained as previously described [40]. Animals were anesthetized in 0.2% (w/v) levamisole prepared in M9 buffer and mounted directly between a glass slide and a coverslip. Imaging was performed at room temperature using a Leica DMI6000B inverted microscope equipped with a 4×/0.10 NA objective lens, a spinning-disk confocal head (Yokogawa CSU10), and an EM-CCD camera (Hamamatsu ImagEM). Image tiling was performed using Metamorph software, and subsequent stitching of the tiles was carried out in ImageJ.

#### Automated multi-worm morphological feature extraction from images

After image acquisition of developmental growth animals, we employed the Segment Anything Model (SAM) [10] to generate automatic mask predictions for each image in our dataset. The ViT-H variant of SAM was utilized, with the following custom parameters: points_per_side: 32, pred_iou_thresh: 0.95, stability_score_thresh: 0.95, min_mask_region_area: 1500. These parameters were empirically determined to optimize mask generation for worm-like objects. The SAM model was implemented using PyTorch [41] and Python 3.10.12 and executed on a CUDA-enabled GPU for efficient processing. For each mask predicted by SAM, we extracted a cutout from the original image. Masks touching image edges were excluded to avoid partial worms.

487 generated cutouts from the developmental growth, first set up acquisitions underwent manual classification by *C. elegans* experts. Each cutout was labeled as either “worm” or “not worm” using the LabelBox web tool with academic access [42]. The classified cutouts were randomly split into training (80%) and validation (20%) sets. We employed transfer learning using a Vision Transformer (ViT-H-14) [43] pre-trained on ImageNet [44] as our base model. The classifier was fine-tuned on our worm/notworm dataset. The “worm” classification included partial worms to improve future usability of the model, although these were later filtered out. The base model’s parameters were frozen, and only the final classification layer was retrained. Data augmentation techniques, including random resized crops and horizontal flips, were applied to the training set. The model was trained for 25 epochs using stochastic gradient descent (SGD) optimization with a learning rate of 0.001 and momentum of 0.9. A step learning rate scheduler was implemented, decaying the learning rate by a factor of 0.1 every 7 epochs. Cross-entropy loss was used as the optimization criterion. Training was conducted on a single GPU using PyTorch. The model achieving the highest validation accuracy (100%) was selected as the final model.

Pre-computed mask predictions from SAM were classified as “worm,” “not worm,” or “background” by the fine-tuned classifier. To address instances of overlapping worm masks (that is, when there were multiple masks produced for the same worm representing subparts of the body), an adjacency matrix was constructed to identify overlapping regions. Connected components analysis was then performed to group overlapping masks. Within each group, only the mask with the largest area was retained for further analysis, ensuring each worm was represented only once. Touching worms were automatically resolved by SAM.

For each retained worm mask, we extracted the following morphological features: area (sum of pixels within the mask), perimeter (Euclidean number of pixels of the shape’s contour), 100-point interpolated medial axis (the longest path along the worm’s centerline identified through graph analysis), length (Euclidean pixel count of the medial axis), width (estimated at each point along the medial axis).

### Droplet swimming recording and analysis

#### Droplet swimming recording set up

Wild-type N2 *C. elegans* were grown at 20 C° and were fed OP50 *E. coli* bacteria as described in Guisnet et al. [39]. A large 10 cm diameter plexiglass dish that was first cleaned with ethanol and a Kimwipe. It was then grounded by bringing the surface in very close proximity to a metal cabinet. This removed static from the plexiglass helping small droplets to stick to the surface. 50 µL of filtered NGML was then pipetted onto the center of the dish. Young adult worms were selected from maintenance plates and left to roam off food on an empty agar plate for 2 minutes before being moved to the droplet with a platinum worm pick.

Uncompressed AVI videos were recorded for 30 seconds at 10 fps with a FLIR Blackfly (BFS-U3-51S5M-C) 5.0 MP camera and Spinnaker SDK software (Teledyne) at 2448 x 2048 resolution. The recording dish was held upside down with a custom made rig and recorded from below. An LED light ring of the same size as the plate was positioned centered with the plate, but further up (∼ 8 centimeters), to minimize light refraction artifacts in the agar and dish plastic.

#### Automated quantitative analysis of swimming behavior

Worm segmentation was performed using the Segment Anything Model 2 (SAM2) [11]. The model was initialized using a pre-trained checkpoint (sam2_hiera_large.pt) and GPU acceleration was employed using CUDA, with TensorFloat-32 compatible hardware. A generic prompt frame was added to all videos and used to automatically guide segmentation on the worm object. Segmentation was propagated backwards from the added prompt frame through the video. To improve segmentation accuracy, a second pass of segmentation was performed on cropped regions of interest. The SAM2 model was applied again to these cropped frames, using a new generic prompt frame matching the crop size. Segmentation was again propagated backwards through the cropped frames. The resulting worm high-definition masks were used for further analysis.

SAM2 masks from the high-definition segmentation were cleaned using morphological operations to remove occasional small particles. Worm skeletons were extracted by mask skeletonization. A custom algorithm was implemented to correct self-touching skeletons, ensuring a single, continuous skeleton for each frame. Skeleton points were smoothed using spline interpolation to 100 points. In addition to those described in the morphological extraction section above, several other metrics were extracted from the skeletons. Gaussian-weighted curvature was calculated along the smoothed skeleton using a window size of 50 points and sigma of 10. Amplitude along the worm’s body was measured as the perpendicular distance from skeleton points to the worm’s centerline (defined as the straight line from head to tail). Worms were classified by shape. Classification was based on curvature and amplitude values. Thresholds for classification were empirically determined. Worms were classified as turned on the z-plane, if their length was smaller by more than 1 standard deviation from the video mean. Wavelength was based on curvature peaks for S-shaped worms, set at 2 worm length for C-shapes, and none for straight or turned worms. Wave number was calculated as the ratio of worm length to wavelength. Spatial frequency analysis was performed using Fast Fourier Transform on the curvature data. Temporal frequencies were calculated from the curvature data. The power spectral density (PSD) of the normalized curvature time series was calculated using Welch’s method with a window size of 30 and an overlap of 25. A sliding window analysis was conducted to identify the dominant frequencies within each segment. Peaks in the PSD were detected, and the frequency corresponding to the peak with the maximum power was selected as the dominant frequency. If no peaks were found, the frequency with the maximum power was used. The dominant frequencies were interpolated across the entire time series to ensure a frequency value for each frame.

### Crawling recording and analysis

#### Crawling recording set up

Wild-type N2 *C. elegans* were grown under standard conditions at 20 C° and were fed OP50 *E. coli* bacteria as described in Guisnet et al. [39]. Young adult worms were selected from maintenance plates and left to roam off food on an empty agar plate for 2 minutes before being recorded. AVI videos were recorded for 1 minute with the same setup and settings as described for swimming.

#### Automated single worm high-definition tracking

Worm tracking was performed in a similar way as described for swimming with a few differences for path tracking and addressing more complicated body postures. For the high-definition segmentation, two generic prompt frames with complicated shapes were added on cropped frames and the SAM2 model was reinitialized and applied to each of the cropped frames one by one. Segmentation was propagated through these short cropped sequences to generate the high-definition masks. The resulting cropped masks were then resized and mapped back to the original frame dimensions for further analysis.

Skeleton morphological features were extracted as described in the swimming section with additional extraction of head bends. Due to the low resolution of the worm in the large image space, head and tail differentiation was not possible by traditional method of brightness difference [28]. A temporal tracking algorithm was employed instead, utilizing a 5 frame sliding window to analyze movement patterns. Endpoint grouping was optimized using the Hungarian algorithm, with the more mobile group designated as the head based on cumulative displacement. Error correction was applied for sudden jumps by cross-referencing with recent history and worm movement. Then, head bending magnitude was quantified by calculating the angle between the head endpoint and the main body axis.

For path analysis, the segmentation masks from previous shape analysis were used to obtain the centroid position for every frame using center-of-mass computation. To reduce measurement noise inherent from changes in the worm’s shape, centroid trajectories were smoothed using Savitzky-Golay filtering, which preserves the underlying movement patterns while eliminating high-frequency noise. Worm movement was classified into three behavioral states: forward locomotion, backward locomotion, and stationary periods. For each frame, instantaneous velocity vectors were calculated from the change in centroid position between consecutive frames. The orientation of the movement was determined by the vector from the body centroid relative to the head position. When centroid direction was towards the head, movement was classified as forward; centroid direction away from the head was classified as backward movement, while low-magnitude velocities (less than 0.5 pixels per frame) were classified as stationary behavior. To ensure accurate movement classification, some corrections were applied. These arose during stationary and turning periods, where head detection was less accurate. Frames with movement classifications inconsistent with surrounding temporal context were identified and corrected. Continuous periods of each movement type were identified as behavioral bouts. For each bout, both temporal duration (in frames) and spatial extent (total distance traveled) were calculated. Movement metrics were calculated for each analyzed video, including, for example, total path length, maximum displacement from origin, proportion of time spent in each behavioral state, and average bout duration for forward and backward movement.

### Calcium imaging in semi-restricted worms

#### Microfluidic imaging of sensory-evoked calcium dynamics in RIA

For recordings of RIA interneuron calcium dynamics, the strain MMH99 (pglr-3a::GCaMP3.3) was used. The worms were grown under standard conditions at 20 C°, fed OP50 *E. coli* bacteria and assayed at the early adult stage [45].

Time-lapse fluorescence imaging of the RIA interneuron in living semi-restrained animals was performed in a microfluidic device as described in [46]. The aforementioned MMH99 strain selectively expressing GCaMP3.3 in RIA were used to measure calcium dynamics within the three distinct axonal compartments of RIA. Streams of liquid alternating between neutral NGM buffer and the attractive odorant IAA (isoamyl alcohol) were passed in front of the animal’s freely moving head to elicit sensory-evoked calcium events [47]. 512 x 512 TIFF stack recordings were acquired for individual animals over 1 minute at 10 frames per seconds and 100 ms exposures on an Olympus IX83 inverted microscope using a 40x U-PlanS Apo (N.A. 1.25) silicone immersion objective, Hammamatsu Orca Flash 4.0LT SCMOS camera, with the cellSens software (Olympus). Automatic stimulus stream switches every 10 seconds were controlled by a solenoid pinch valve array (Automate Scientific).

#### Manual segmentation of axonal compartments and head bending extraction

The brightness of the RIA compartments and the worm’s head position were manually extracted for comparison using the Fiji software as described in Ouellette et al. [23].

#### Automated segmentation of axonal compartments and head bending extraction

Full-frame videos were processed to extract a region of interest centered on the RIA interneuron. Like for the other modalities, the Segment Anything Model 2 (SAM2) video predictor was initialized using the Hiera Large checkpoint. For each video, a single annotation point was placed on the RIA region in the final frame. This annotation was propagated backwards through all frames. The center of mass for each RIA mask was calculated across all frames to create a fixed crop window of 110×110 pixels around the RIA region throughout the video sequence. To simultaneously segment the three anatomical structures within the cropped RIA region (nerve ring dorsal (nrD), nerve ring ventral (nrV), and loop. A prompt management system was developed to enable consistent segmentation across videos. Expertly annotated prompt frames were incorporated into the video sequences to guide the segmentation model. SAM2 was reinitialized with the augmented video sequences, and structure-specific prompts. A comprehensive quality analysis was performed to identify segmentation artifacts, including empty masks, oversized regions, and overlapping structures. Additional prompt frames were added to the prompt pool if refinement was necessary. On rare occasions, sporadic frames with a missing mask were filled by interpolation. Occasional small noise artifacts were removed through connected component analysis. Brightness measurements were extracted from the segmentation masks by extracting the pixel intensities from the corresponding grayscale images. Background correction was performed by sampling 100 background regions located at least 40 pixels away from any segmented object, as determined by distance transform analysis. The mean background intensity was subtracted from object measurements to obtain corrected brightness values. Since animals were positioned either on their left or right side in the chip, the spatial relationship between the loop structure and nrD was analyzed to adjust head bending angles as dorsal or ventral.

Full-body segmentation was performed on the original full-frame videos to enable postural analysis. A single annotation point was placed on the organism body and propagated through all frames using SAM2. The resulting masks captured the complete body, which served as input for subsequent head angle analysis. Morphological skeletonization was applied to whole-body masks to extract one-pixel-wide centerlines representing the organism’s medial axis. The angle between head and body vectors (anterior and posterior portions of the skeleton) was computed for each frame. Temporal smoothing was applied to remove measurement noise while preserving genuine behavioral transitions. A small temporal sliding window smoothing of 3 frames was applied to remove noise from small variations in body segmentation outline. Final head angle sign was adjusted based on the organism’s orientation determined in the previous step.

### Hardware and code

All image analysis, model training and model inference was done on a local workstation with Ubuntu 22.04 and one Nvidia RTX 3090 GPU. To speed up processing, some operations were scaled to 48 parallel cores. For image analysis, the Meta SAM model sam_vit_h_4b8939 was used. For video analysis, the Meta SAM2 repository was cloned and the model downloaded from the July 28th 2024 version (https://github.com/facebookresearch/sam2/). All other data analyses and visualizations were conducted using Python (v3.12.3), uv for environment management and the following openly accessible libraries: h5py, matplotlib, numpy, OpenCV, pandas, PyTorch, scikit-image, scikit-learn, scipy, seaborn and tqdm [41,48–59]. All code is freely accessible at https://github.com/lillyguisnet/TWARDISv0.1. Our fine-tuned worm classifier is available on HuggingFace (https://huggingface.co/lillyguisnet/celegans-classifier-vit-h-14-finetuned).

## Acknowledgements

We thank Laeya Baldini and Stephanie C. Weber from McGill University for sharing their developmental growth images used as the second dataset in this publication. Maxime Rivest for guiding early readings on computer vision and troubleshooting workstation issues. Meta FAIR for releasing the SAM models open-source. All members of the lab for helpful discussions and comments. Strains were provided by the *Caenorhabditis* Genetics Center (CGC), which is funded by NIH Office of Research Infrastructure Programs (P40 OD010440).

## Supporting information

**S1 Fig. Scatterplot of Fiji- versus TWARDIS-estimated head angles with sign ignored.** Fiji estimations (−1, 1) are converted to the (−90, 90) range for comparison. 0 represents a perfectly straight head. +/− 90 degrees represent maximal bending. r = 0.746. n = 13,486 frames from 22 recordings.

**S1 Movie. Droplet swimming recording with worm mask and body curvature kymograph.** Background image of original recording with resolution halved. Picture-in-picture in top left corner shows the final worm mask for every frame. Bottom live plot shows Fig 2E, the gaussian-weighted curvature intensity of 100 interpolated points along the worm’s body per frame (same legend as Fig 2E).

**S2 Movie. Crawling recording with worm mask, head angle and speed.** Background image of original recording with resolution halved. Picture-in-picture in top left corner shows the final worm mask for every frame. Dots of the worm position are overlaid on the original frames to represent the worm trace through time. Top live plot shows Fig 3E: the head bending angle relative to body in degrees per frame where 0 indicates a straight head relative to the body, and positive and negative values represent bends on either side, but ventral and dorsal sides are not distinguished, and points along the trace represent the detected peaks and troughs used for head bend metrics. Bottom live plot shows Fig 3F: the worm speed by frame, where positive values represent forward motion, negative values represent backward motion and stationary motion is determined by absolute speed less than 0.6 pixel/second (shaded area). This movement classification is also color-coded as in Fig 3C for the dot overlays and for every frame at the bottom left of the original image.

**S3 Movie. RIA calcium imaging recording with axonal compartment segmentation, head angle, and calcium activity.** Background image of original recording with no modification. Final body segmentation is overlaid on the original frames in purple for every frame. Head angle in degrees extracted from body segmentation is presented for every frame over the head of the worm and in the top-most live plot (same as Fig 4G). Boxed picture-in-picture shows the final masks at every frame for each segmented axonal compartment: nrD (orange), nrV (blue) and loop (green). Resulting normalized fluorescence intensity traces for each compartment are shown in the bottom three live plots.

**S1 Dataset. Human and TWARDIS-extracted area and perimeter in pixels.**

**S2 Dataset. Time-series metrics of a 10 frames-per-second one minute recording of a wild-type worm swimming in a liquid droplet.**

**S3 Dataset. Time-series metrics of a 10 frames-per-second one minute recording of a wild-type worm crawling on an agar plate with food.**

**S4 Dataset. Fiji- and TWARDIS-extracted head bending and fluorescence time-series values of 10 frames-per-second one minute recordings of wild-type worms in a microfluidic device expressing GCaMP3.3 in RIA.**

**S5 Dataset. Normalized fluorescence intensity and average absolute difference from mean for Fiji- and TWARDIS-extracted data of the loop compartment.**

**S6 Dataset. Fiji- and TWARDIS-estimated head angles centered normalized from 0 to 180 degrees.**

